# Age-related changes in microRNAs expression in cruciate ligaments of wild-stock house mice

**DOI:** 10.1101/2021.12.01.470740

**Authors:** Yalda A. Kharaz, Katarzyna Goljanek-Whysall, Gareth Nye, Jane Hurst, Anne McArdle, Eithne J. Comerford

## Abstract

**Aim:** Cruciate ligaments (CLs) of the knee joint are commonly injured following trauma or ageing. MicroRNAs (miRs) are potential therapeutic targets in musculoskeletal disorders. This study aimed to 1) identify if wild-stock house (WSH) mice are an appropriate model to study age-related changes of the knee joint and 2) investigate expression of miRs in ageing murine CLs.

**Methods:** Knee joints were collected from 6 and 24 months old C57BL/6 and WSH mice (*Mus musculus domesticus*) for histological analysis. RNA extraction and qPCR gene expression were performed on CLs in 6, 12, 24, and 30 month WSH old mice. Expression of miR targets in CLs was determined, followed by analysis of predicted mRNA target genes and Ingenuity Pathway Analysis.

**Results:** Higher CL and knee OARSI histological scores were found in 24 month old WSH mice compared to 6 and 12 month old C57BL/6 and 6 month old WSH mice (*p*< 0.05). miR-29a and miR-34a were upregulated in 30 month-old WSH mice in comparison to 6, 12 and 24-month-old WSH mice (*p*<0.05). Ingenuity Pathway Analysis on miR-29a and 34a targets was associated with inflammation through interleukins, TGFβ and Notch genes and p53 signalling. Collagen type I alpha 1 chain (COL1A1) correlated negatively with both miR-29a (r= -0.35) and miR-34a (r= -0.33).

**Conclusion:** The findings of this study support WSH house mice as an accelerated ageing model of the murine knee joint. This study also indicated that miR-29a and 34a may be important regulators of COL1A1 gene expression in murine CLs.

## Introduction

The anterior and posterior cruciate ligaments are the main stabilisers of the knee joint [1]. Their composition consists of fibroblasts embedded in a specialised extracellular matrix (ECM), comprised mainly of type I collagen, and elastin with a range of non-collagenous proteins and proteoglycans [2]. The knee joint anterior cruciate ligament (ACL) is one of the most frequently injured ligaments [3] resulting in significant joint instability, immobility [3], muscle atrophy [4] and induction of knee joint osteoarthritis (OA) [5]. Knee joint OA has major physical, social and financial implication for the ageing population [6]. ACL injury caused by trauma or contact sport only accounts for about 30 percent of ACL injuries [7]. The remaining ACL tears are from non-contact injuries, which can occur following gradual degeneration of the ligament extracellular matrix (ECM) [7]. To date the exact etiopathogenesis of non-contact ACL ruptures is not defined, however risk factors have been identified such as age [8], gender [9], bodyweight [10] and genetics [11]. The healing potential of the ACL is poor and reconstruction following traumatic injuries does not completely restore the functional stability of the knee joint [12], which contributes to development of knee osteoarthritis [13]. Therefore more effective novel strategies for managing ACL injury should be developed to promote the healing of the native ligament structure to have a protective impact on the tissue mechanics and subsequently the articular cartilage of the knee [14].

Ageing has been shown to alter ligament extracellular matrix composition [8, 15-17]. This has been demonstrated in aged human and canine ACL, where changes in the ECM ultrastructure have been reported prior to cartilage injury and other signs of knee joint osteoarthritis [8, 15]. Amiel et al.(1991) [16] demonstrated that collagen content and synthesis decreased with age in knee ligaments from rabbits resulting in significant effect upon the mechanical properties of these tissues. ACLs from healthy aged dogs (and dog breeds at a high risk to ACL injury) show increased chondroid metaplasia of ligament fibroblasts and reduced cell density compared with ACLs from young and low risk animal [17]. Similar changes in degenerated and ACLs of aged human occur with decreased fibroblast density being found in ACLs of older people [15]. To date, there are no current treatment options targeting the prevention of ligament ECM degradation leading to ligament damage and eventual rupture in high disease risk species such as dogs or man.

MicroRNAs (miRNAs or miRs) control the simultaneous expression of many genes and have been proposed as potential therapeutic molecules for disorders of musculoskeletal tissues such as ligament due to their relatively easy delivery into tissues [18]. They are small endogenous (∼22□nt) noncoding RNAs that play important regulatory roles in animals and plants through posttranscriptional modulation of gene expression by binding and repressing the expression of specific mRNAs [19]. miRNAs have been demonstrated to play a role in disease repair mechanisms in a number of different tissues, including tendon and ligament [14, 20, 21]. A recent study in human ACLs has demonstrated differences 39 differentially expressed miRs between patients with and without knee OA. Twenty-two miRs such as 26b-5p and 146a-5p were found to be upregulated, while 17 miRs such as 18a-3p and 138-5p were downregulated in the osteoarthritic ACL tissues, suggesting that related miRNA dysregulation is involved in ligament injury in patients with OA [21]. Another study demonstrated that miR-210 was decreased in partially transected ACLs in a rat model suggesting that miR-210 promotes ACL healing through enhancement of angiogenesis [14]. To date, there is little known on the role of miRNAs and their expression in ACLs during ageing and limited data on their use in either treating or preventing ligament injuries.

There are currently numerous rodent models that are used in orthopaedic research, including mice, rats, gerbils and squirrels, among others; with mice and rats being the more frequently used models [22, 23]. Laboratory mice such as the C57BL/6 strain have been domesticated, possibly over a period of up to 3000 years [24] and show considerably reduced activity and speed of movement compared with wild house mice due to both genetic changes and substantial environmental constraints [25]. In comparison to C57BL/6 mice, an inbred strain of house mice *(Mus musculus domesticus)* derived from the wild in 1978 has been found to exhibit considerably greater activity under standard caged conditions [26], suggesting that laboratory animals more recently derived from the wild may provide a more appropriate model to study age related changes associated with normal physical activity. In this study we hypothesised that 1) wild-stock mice are an appropriate model in comparison to C57BL/6 mice to study age-related changes in the knee joint associated with normal day-to-day activities and 2) there are age-related changes in miRNAs expression in cruciate ligaments of mice associated with normal physical activity. Therefore, this study aimed to compare age-related changes between wild-stock house mice and C57BL/6 using histological analysis, and to investigate miRNAs expression in cruciate ligaments during ageing of mice associated with normal physical activity.

## Methods

### Animals

Six and 24 months old Specific Pathogen Free (SPF) C57BL/6 mice were purchased from Charles River (Lyon, France) and delivered to the Biomedical Services Unit at the University of Liverpool at least 1 month prior to the required age to allow acclimation. Mice were individually housed and were fed a CRM (P) rodent diet with ad libitum access to food and were maintained under barrier conditions in microisolator cages on a 12-h dark/light cycle. Procedures were performed in accordance with UK Home office Guidelines under the UK Animal (Scientific Procedures Act 1986) and received ethical approval from the University of Liverpool Animal Welfare Committee.

Wild-stock house mice (*Mus musculus domesticus*) were originally derived from local Cheshire populations, outbred in captivity for 1-4 generations [27]. Animal use and care was in accordance with EU directive 2010/63/EU and UK Home Office code of practice for the housing and care of animals bred, supplied or used for scientific purposes. The University of Liverpool Animal Welfare Committee approved the maintenance of our wild mouse colony, but no specific licenses were required. These mice were euthanized for reasons unrelated to this study and the knee joint tissue was obtained post mortem as clinical waste. Mice were fed with Corn Cob Absorb 10/14 substrate (IPS Product Supplies Ltd, London, UK) and with *ad libitum* access to food (LabDiet 5002, Purina Mills) and water. All wild-stock mice were provided with paper wool nesting material (IPS Supplies Ltd) and a variety of cardboard tubes or boxes, and plastic tube or clip-on shelter hanging from the cage top. Subjects were maintained under controlled environmental conditions: temperature 20–21 °C, relative humidity 45–65 % and a reversed 12:12 h light cycle.

### Tissue collection

The entire knee joint from 6 and 24 month old C57BL/6 and wild-stock house mice (n=6) were collected and stored in 4% paraformaldehyde for 48 hours for histological analysis. For RNA extraction, ACLs and PCLs were isolated with a dissecting microscope (Olympus CK40) from 6, 12, 24 and 30 month old wild stock house mice, snap frozen and stored at - 80°C until required.

### Knee joint collection and histological analysis

Following fixation, knee joints were decalcified for four weeks in a solution of 25 g EDTA in 175 cm^3^ distilled water (pH= 4-4.5), wax-embedded and 6 μm coronal sections cut through the whole joint [28]. Sections were scored twice by one observer blinded to the sample origins using the OARSI grading system for rat knee joint cartilage [29] and using a comparative grading system for ligament scoring. Both ACLs and PCLs were scored based on strength of ECM staining, cell hypertrophy, cell clustering, loss of alignment and ossification and were graded from 0-4 based on the extent of changes ((0= normal 0% increased; 1=5-25% increase mild abnormality; 2= moderate abnormality, 26-50% increase; 3= marked abnormality, 51-75% increase; 4= severe abnormality, 76-100% increase) [2].

### MicroRNA: target interaction prediction

Experimentally predicted miRNAs from previously identified ECM proteins/gene in ligament [30] was determined using Targetscan Human (Version 7.1) as described previously [18] (Table 1). MicroRNAs predicted to regulate several ECM components of ligament were investigated in order to identify miRs that regulate ligament programmes, pathways and networks rather than individual genes.

**Table 1.**
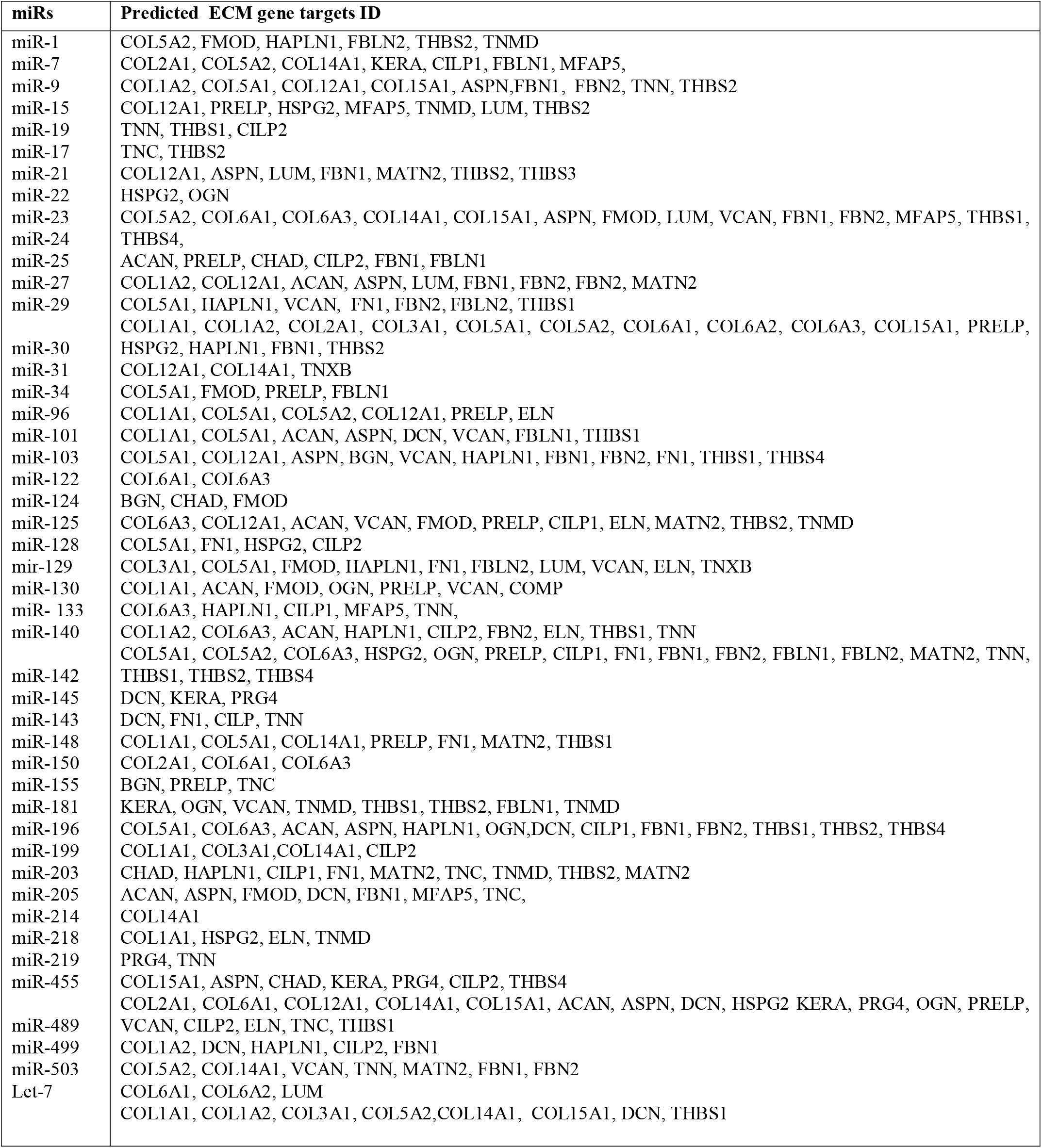
Identification of predicted biological targets of microRNA (miR) in murine cruciate ligaments through TargetScan.

### Interaction network analysis

Pathway analysis of differentially expressed miRNAs was produced using QIAGEN Ingenuity Pathway Analysis (IPA) Product (Ingenuity_Systems, http://www.ingenuity.com), the Core Analysis function and the Path Designer feature [31]. Network interaction maps of predicted ECM target genes were created by the differently expressed miRNA using String bioinformatics tool (String-DB) version 9.1 by allowing for experimental evidence in addition to the predicted functional links: co-occurrence, co-expression, databases, and text-mining [32].

### RNA isolation and Real Time PCR

Total RNA isolation was performed using standard methods as described previously [33]. Details of primers are included in Supplementary Tables 1 and 2. In brief, RNA was extracted with Trizol (Invitrogen^™^ Life Technologies, USA), quantified according to the manufacturer’s protocol using a Nanodrop ND-100 spectrophotometer (Labtech, Uckfield, East Sussex, UK) and assessed for purity by UV absorbance measurements at 260 and 280 nm. cDNA synthesis for miRNA was performed using 200ng RNA and miRscript RT kit II (Qiagen, UK) according to the manufacturer’s protocol. MicroRNA qPCR analysis was performed using miRScript Sybr-Green Mastermix (Qiagen, UK) in a 20 µl reaction Applied Biosystems 7500 Fast Real-Time PCR System following the manufacturer’s protocols. The qPCR conditions were: 95°C for 15 mins for initial activation followed by 40 cycles of 95°C 30 seconds, 55°C 30 seconds, 70°C 30 seconds.

cDNA synthesis for gene expression analyses was performed using 500 ng RNA using Moloney murine leukemia virus (M-MLV) reverse transcriptase and random hexamer oligonucleotide (both from Promega, Southampton, UK) using 500 ng RNA in a 25 µl reaction. mRNA qPCR was performed on 5 µl 10x diluted CDNA by employing a final concentration of 300 nM of each primer in 20 µl reaction on ABI 7700 sequence detector using MESA Blue SYBR Green reagent (Eurogentec, Belgium) using the following protocol: denaturation at 95 °C for 5 minutes, followed by 40 cycles of DNA amplification (15 seconds 95 °C and 45 seconds annealing at 60 °C) [34]. Used miRNA primers in this study has been validated in previous publications [18] and supplied by Eurogentec (Supplementary Table 1). mRNA and miRNA data sets were compared with the designated control utilising housekeeping genes *GAPDH* or *RnU6* as detailed in Figure legends.

### Statistical analysis

Statistical analysis was performed on both histological scoring and qRT-PCR using Graphpad Prism (Version 7, GraphPad Software, USA) and a significance level of 5%. The normal distribution for each data set was assessed using a Kolmogorov-Smirnov test. One-way ANOVA with Tukey *post-hoc* test comparing histological scoring between C57BL/6 and wild-stock house mice.

Intra-observer agreement of histological scoring systems was calculated using Cohen’s kappa coefficient (www.statstodo.com/CohenKappa_Pgm.phpl). One-way ANOVAs with Tukey *post-hoc* test was assessed for differences in qRT-PCR data between age groups. Pearson’s correlations (*r*) assessed relationships between differential expressed miRNA and mRNA target gene.

## Results

### Morphological differences between wild-stock house and C57BL/6 mice

An overall Kappa statistic of 0.8 for blinded intra-observer agreement for the histological grading was calculated, indicating strong agreement. Histological changes between 6 (Figure 1A-C) and 24 month C57BL/6 (Figure 1D-F) and 6 (G-I) and 24 month old wild-stock house (J-L) showed only ultrastructural abnormalities in CLs of the 24 month old wild-stock house mice with chondrocytic cell morphology and increased proteoglycan content around cells evident (Figure 1J and K black arrows). Twenty-four-month-old wild-stock house mice also showed similar abnormal ultrastructural changes in both CLs with severe cartilage lesions extending >75% of the articular surface being observed (Figure 1L, black arrows). In addition, osteophyte formation and structural changes in meniscus were also observed around and in the knee joints respectively of the 24 month old wild-stock house mice (Figure 1K, white arrows). No effect of age was seen in CL (Figure 1M) or and OARSI scores (Figure 1N amd O). In contrast both the CL (Figure 1M) and OARSI mean and maximum scores (Figure 1N and O) was significantly higher in 24 month old wild-stock house mice (*P*<0.05 in comparison to 6 month old wild-stock house mice as well as 6 month and 24 months old C57BL/6 mice.

**Figure 1.**
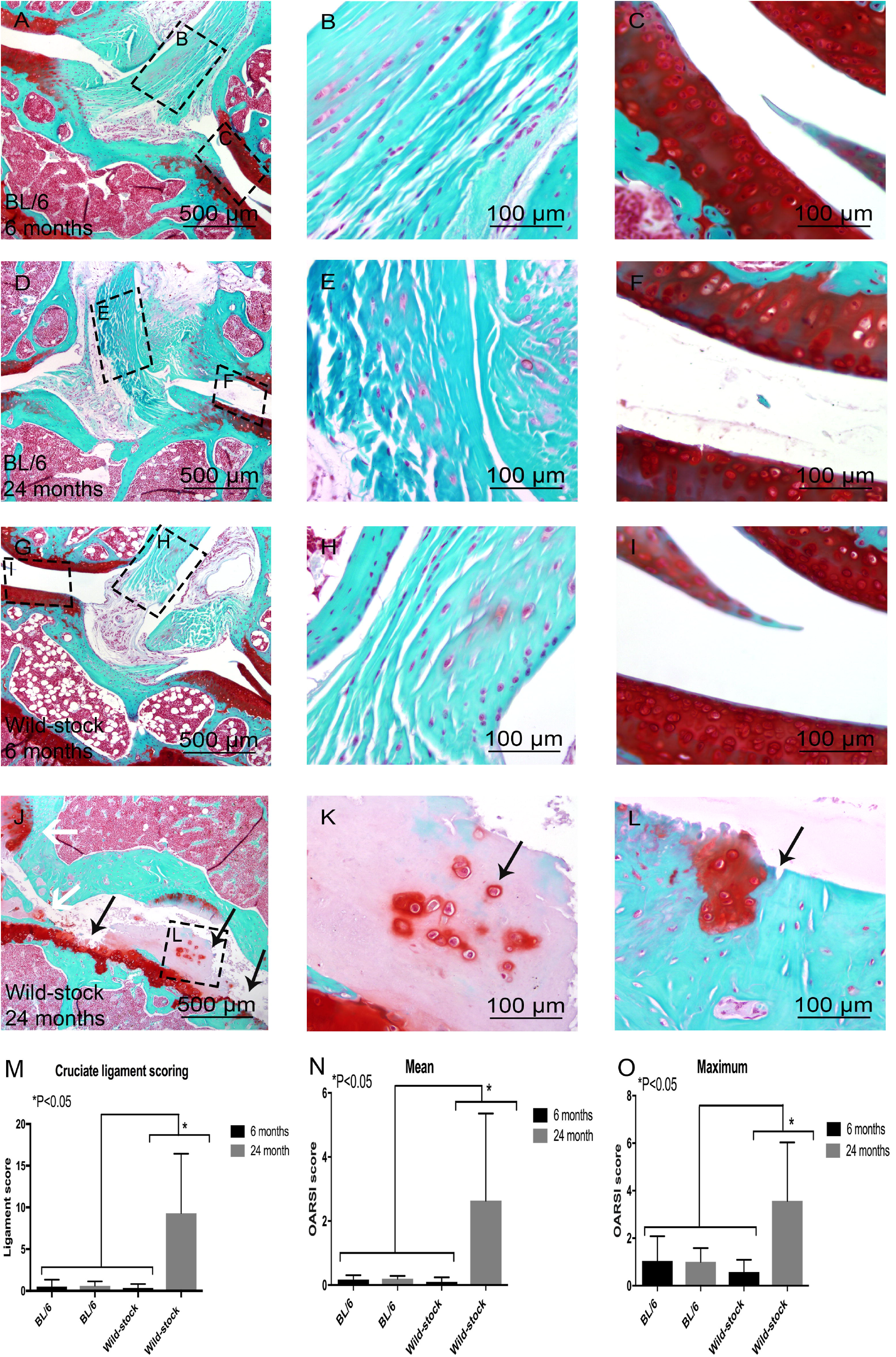
Histological comparison between 6 month old C57BL/6 (A-C), 6 month old wild-stock house mouse (D-F), 24 month old C57BL/6 (G-I) and 24 month old wild-stock house (J-K) mouse with Safranin-O knee joints. An abnormal structure of cruciate ligaments with chondrocytic cell morphology and increased proteoglycan content around cells was observed (black arrow in J and K). Erosion to the calcified cartilage extending >75% of the articular surface (black arrow in J and L) and osteophyte formation and structural changes in meniscus was also observed (white arrows in J). Statistically significantly higher cruciate ligament and OARSI scores were measured in 24 month old wild-stock house mice compared to 6, 12 month old C57/BL6 and 6 month old wild-stock house mice (M-O). Data are means ± SEM.

### MicroRNA expression and Ingenuity Pathway Analysis

The miRNA expression was only performed in 6, 12, 24 and 30 months old wild-stock house mice. The expression of several miRNAs that were predicted through Target Scan (Table 1) to regulate key ECM components associated with ligament ageing was determined. There were not any significant differences in expression level between any of the age groups with miR-128, miR-455, miR-143, miR-21, miR-34a and miR-181 (Figure 2). However, miR-29a and miR-34a were expressed at significantly higher levels in 30 month-old mice (*P*<0.05) in comparison to the 6 month and 24 old mice (Figure 2). Ingenuity Pathway Analysis indicated that upregulation of miR-29a and miR-34a during ageing in CLs may be associated with MAPK (mitogen activated protein kinase), p53, SMAD and Notch signalling, as well inflammation-related genes such as TGFβ (transforming growth factor beta) and interleukin-2 (Figure 3A). String-DB analyses of ECM-associated predicted target genes of miR-29a and miR-34a revealed a network of genes containing one highly connected cluster around collagens ECM organisation (Figure 3B).

**Figure 2.**
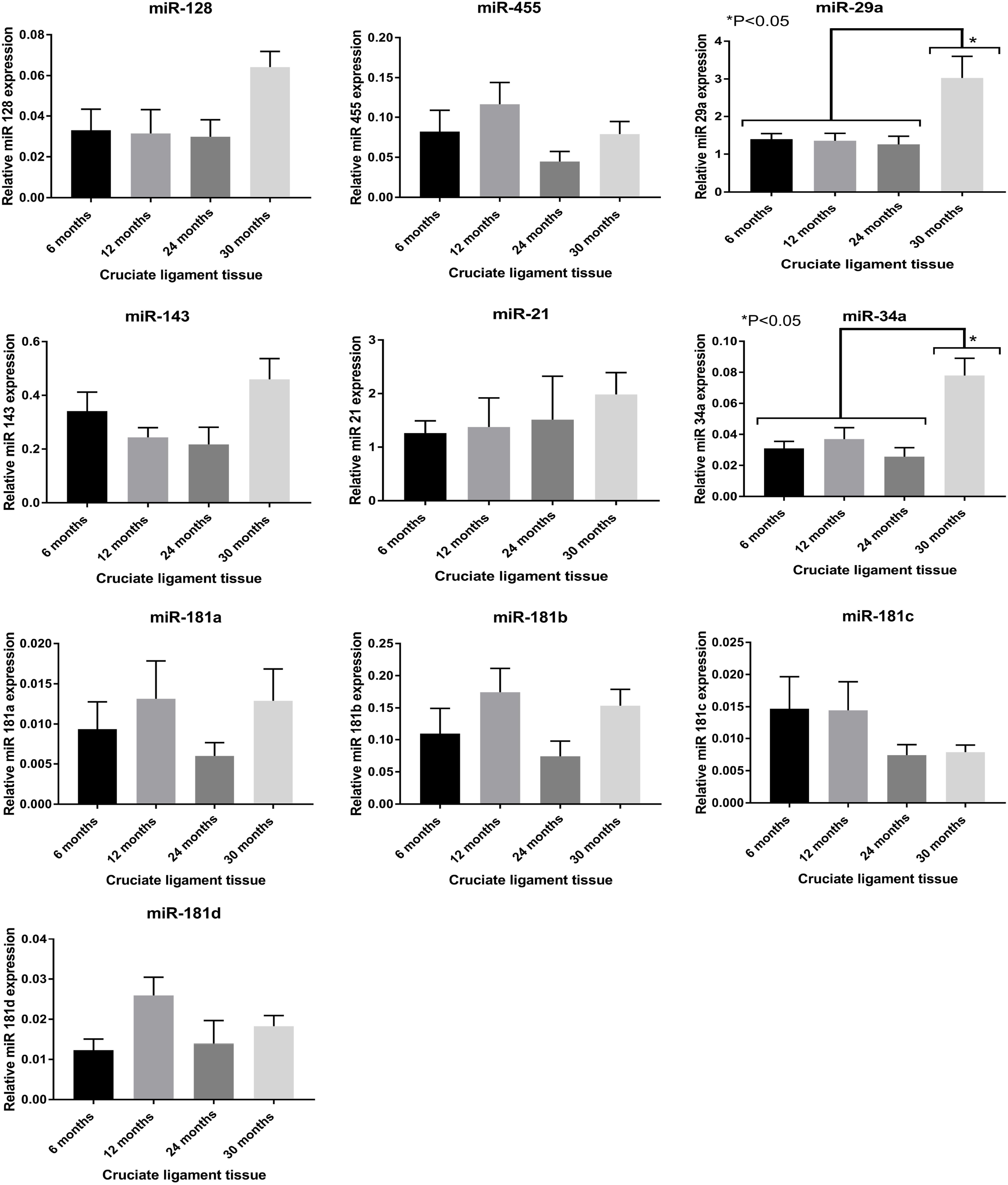
MicroRNA (miR) relative expression (to housekeeping gene *Rnu6*) at different ages in murine cruciate ligaments (data are means ± SEM). Statistically significant higher expression of miR-29a and miR-34a was found in 30 month old mice in comparison to the 6, 12 and 24 month old mice. Data are means ± SEM.

**Figure 3.**
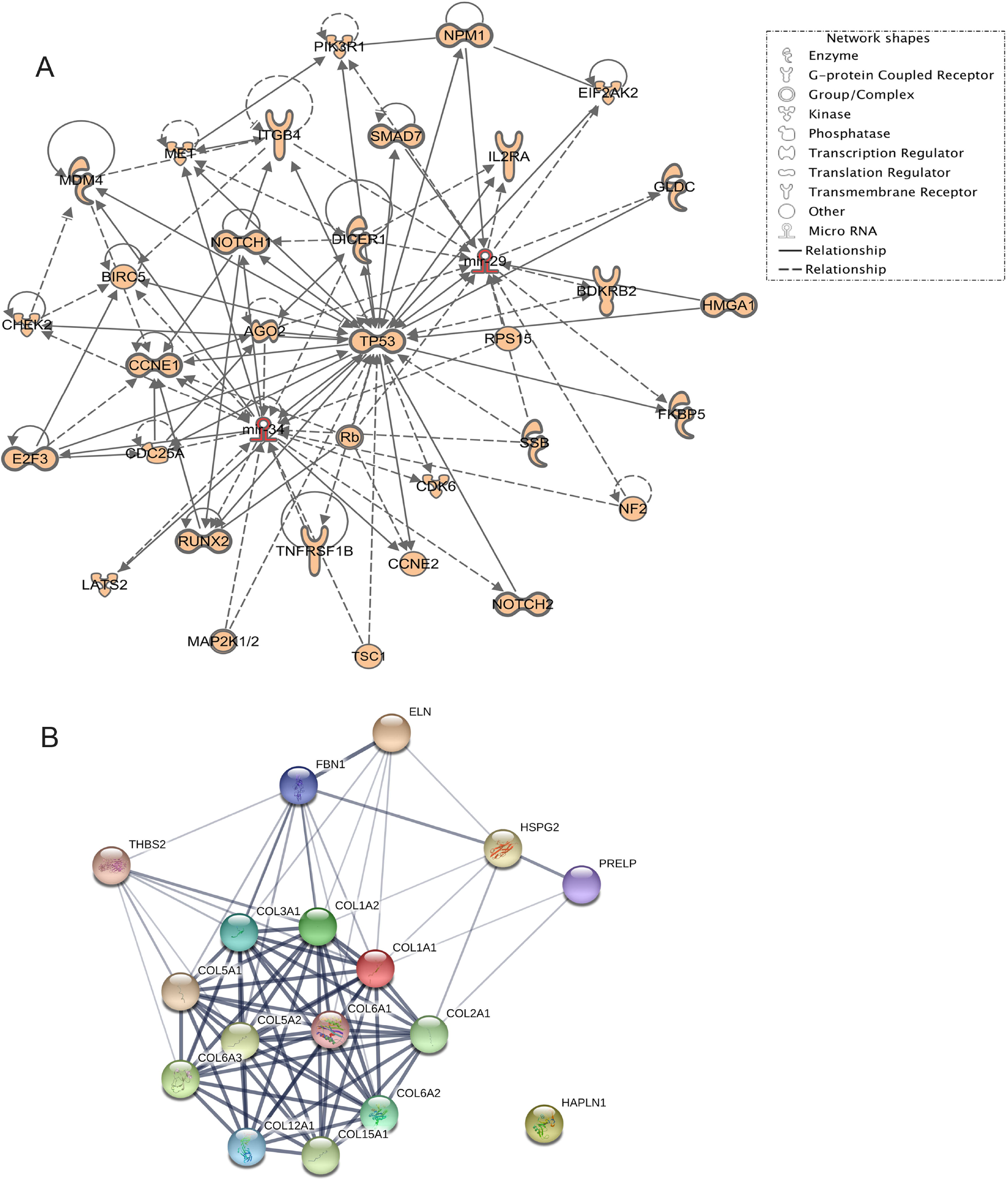
(A) Network of interactions constructed for the microRNAs (miR-29a and miR-34a) targets using Ingenuity Pathway Analysis software (IPA). (B) An interaction map of extracellular matrix target genes predicted through Target Scan to be upregulated by both miR-29a and miR-34a was built with STRING. One highly connected cluster was evident around collagen proteins and a high confidence level (0.0700) was allowed for experimentally predicted gene-gene interaction.

### Gene expression of target genes

Target gene expression of several ECM-associated targets predicted to be regulated for both miR-29a and miR-34a using TargetScan and Ingenuity Pathway Analyses was measured. These mRNA targets were also identified previously in non-diseased canine ACLs as ligament markers [30]. Measurement of mRNA targets included collagen type I alpha1-chain (COL1A1), collagen type 3 alpha-1 chain (COL3A1), collagen type V alpha-1 chain, collagen type XII alpha 1-chain (COL12A1) and proline and arginine rich end leucine rich repeat protein (PRELP) (Figure 4A). The expression of COL1A1 was significantly higher in CLs of mice at 12 month (*P*<0.05) as compared to 6, 24 and 30 month old mice (Figure 4A). Significantly higher expression of COL3A1 (*P*<0.05) was also measured in 6 month old mice than 12, 24 and 30 month old mice (Figure 4A). Pearson’s correlation analyses demonstrated statistically significant negative correlations between miR-29a and miR-34a and COL1A1 expression levels (r= -0.41 and r= -0.4) (*P*<0.05) (Figure 4B). Pearson’s correlation analyses of the expression of miR-29a and miR-34a with COL3A1 (r=-0.03 and r=-0.11) was not statistically significant (Figure 4B).

**Figure 4.**
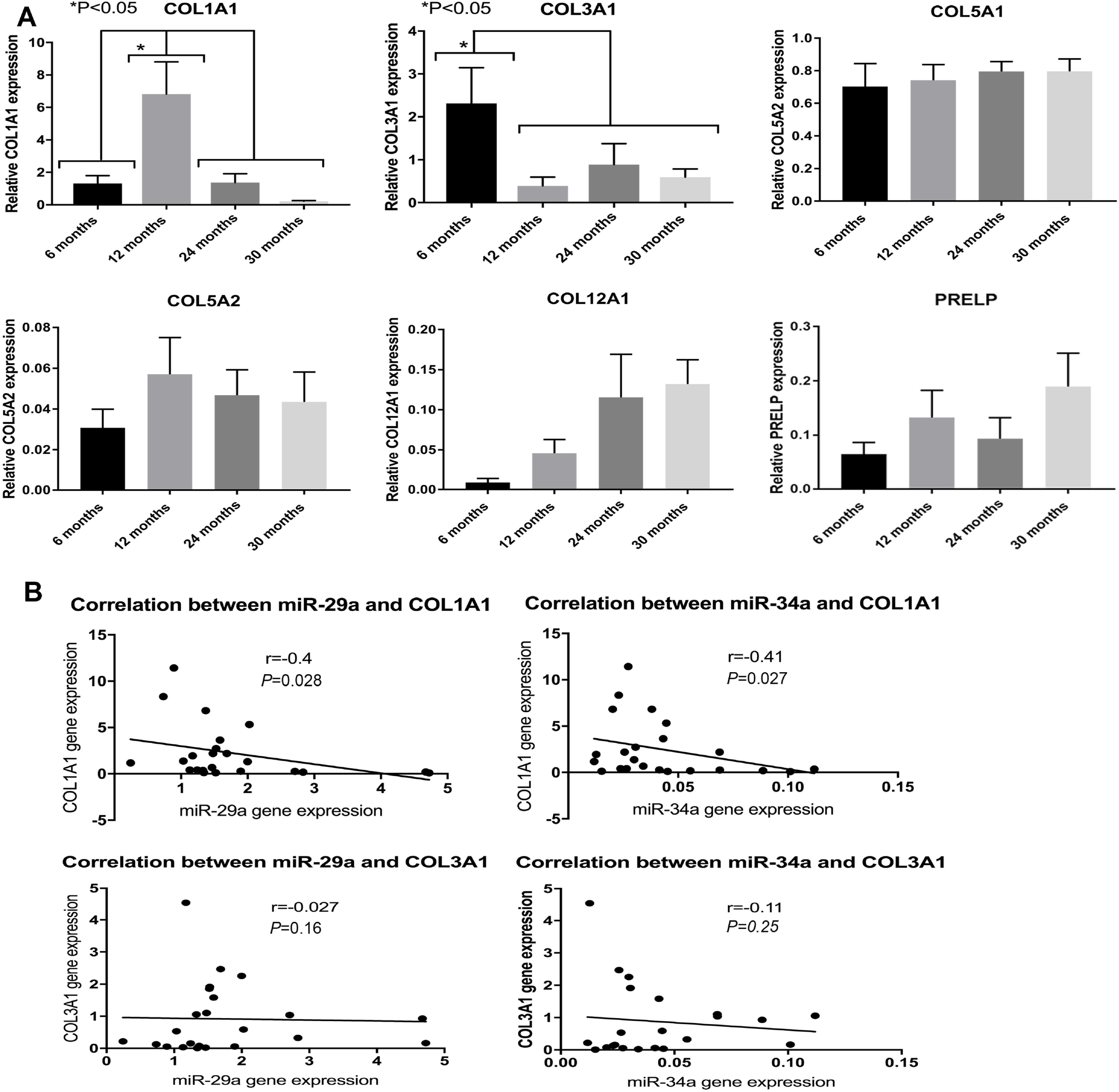
: A. Relative expression (to housekeeping gene *GAPDH)* of extracellular matrix predicted mRNA target genes for miR-29a and miR-34a in mice cruciate ligament. (B) Pearson’s correlation demonstrated statistically significant negative correlations between miR-29a and miR-34 with COL1A1 target gene (r=-0.41, *P*<0.05 and r=-0.4, *P*<0.05). Pearson’s correlation between miR-29a and miR-34a with COL3A1 was not found to be statistically significant. Data are means ± SEM.

## Discussion

This study has identified age-related morphological differences in the knee joints and CL from wild-stock house mice that exhibit normal physical activity, in contrast to no effect age seen in joints and ligaments of C57Bl/6 mice. Wild-stock house mice may therefore be a more appropriate model of healthy ageing compared to joints and ligaments of C57BL/6 mice. This is the first study to measure the differential microRNA gene expression in murine cruciate ligaments with age. Our findings demonstrate that the miRNAs (miR-29a and miR-34a) were differentially expressed in cruciate ligaments of 30 month old-wild-stock house mice. Both miR-29a and miR-34a negatively correlated with COL1A1 gene expression, suggestive of their role as key regulators of the ECM of murine CLs during ageing.

Our histological analysis of the knee joints in different age groups of C57BL/6 and wild-stock house mice demonstrated morphological changes in tissue structures and significantly higher histology scores in knee joint femoral and tibial cartilage, formation of osteophytes and changes in CL structure in the aged wild-stock mice compared with 6 months old wild-stock house mice and age matched C57Bl/6 mice. These findings suggest that wild-stock house mice are a more appropriate model to study for accelerated ligament degeneration and were further used to investigate the regulation of miRNA in ageing CL.s

Amongst a subset of miRs, miR-29a and miR-34a were found to have a significantly higher expression in 30 month old mice in comparison to the other age groups of wild-stock house mice. Both miR-29a and miR-34a have been found to regulate age-related diseases such as vascular ageing [35] and cellular senescence in muscle [36]. Studies into OA pathogenesis in human and mouse cartilage have also demonstrated upregulation of both miR-29a [37] and miR-34 [38]. In the current study, the differential expression of miRs measured in the ageing murine CLs may be associated with the knee joint cartilage erosion and OA pathogenesis of these ageing mice, as observed through our histological staining. Together, these findings may indicate that the upregulation of miR-29a and miR-34a in ageing murine CLs could be associated with late stage pathology of OA and may be key to later stages of joint degeneration.

During ageing, signalling pathways including TGFβ, NOTCH, pSMAD, IGF, MAPK are known to play a role in tissues such as mouse and human skeletal muscle [39, 40] and knee joint cartilage [41]. In the ACL, the exact signalling pathway mechanism is yet to be elucidated during ageing and disease. However, our constructed network using IPA contained signalling pathways such as TGFβ and MAPK, p53, miR-29a and miR-34a. This finding agrees with a previous human OA study where miR-34 affected chondrocyte apoptosis and proliferation by targeting p53 regulatory pathway and silent information regulator 1 during disease pathogenesis [38]. miR-29 has also been found to activate p53 in murine fibroblasts through repression of Ppm1d phosphatase during ageing and DNA damage activation [42]. Together, these data demonstrate the key signalling networks with mice cruciate ageing and association of miRs. Further functional investigation is required to study the association networks between miRs regulation and signalling pathways during cruciate ligament ageing.

In the current study, both miR-29a and miR-34a were predicted to target several collagens genes including type I, II, III, V, VI, XII and other ECM genes such as the PRELP, HAPLN1, HSPG2, FBN1, THBS2. We primarily focused on validation of several target genes that have previously been identified as ligament ECM markers [2, 30, 43] and particularly on target genes COL1A1 and COL3A1 genes, where higher mRNA levels have been found in ruptured compared to normal ACLs [44, 45]. We found significant alterations in mRNA expression levels of collagen types I and III during ageing of murine CLs. The reduced expression of COL3A1 levels in the current study in mice after 6 months of age does not agree with studies of equine superficial digital flexor tendon, where higher levels of COL3A1 were found with ageing [46]. This may suggest differences between COL3A1 turn-over rates between species and tissue types. The increase expression of COL1A1 levels at 6 month of age followed by decreased expression in ageing mice may be due to dynamic interactions and feedback mechanisms between miRNA and targets genes. Apart from miR-29a and miR-34a, other miRs such as miR-133a [47] and miR-129 [48] have also been demonstrated to regulate COL1A1 gene expression in cardiac and liver tissues, but may also be involved in the ageing cruciate ligament in the current study. The decreased expression of COL1A1 after in mice 12 month of age in our study may also have contributed to the abnormal structure of the cruciate ligaments found in our histological analysis. Previous studies have demonstrated changes in ageing equine tendons in their collagen cross-link profile following glycation [49] and an increase in collagen degradation markers [49], which may also agree with the abnormal ECM structure and the decreased levels of COL1A1 in the ageing mice cruciate ligament in this study. Further studies are required to measure accumulation of damage in mice cruciate ligament collagen during ageing.

We found negative correlations primarily between COL1A1 gene expression and miR-29a and miR-34a, suggestive of a potential regulation of this vital gene in cruciate ligament tissue pathology during ageing. In tendon injuries, miR-29a has been demonstrated as a post-transcriptional regulator of collagen type III gene expression in murine and human tendon injury, through interleukin-L33 [20]. However, we did not find any statistically significant negative correlations between either miR-29a and miR-34 and COL3A1, suggesting that miR-29a may play different roles in the ageing versus injury process. Future studies will involve *in vitro* miRNA target validation and *in vivo* delivery of miR-29a and miR-34a in wild-stock mice knee joints followed by phenotypic analyses.

In conclusion, CL ageing and degeneration, resulting in injury can have severe physical, social, economic consequences to the affected individual and will lead to development of degenerative joint diseases such as OA. Through our histological analysis we have shown that wild-stock house mice are an appropriate mouse model to study age-related changes in the knee joint compared C57BL/6 mice. This study also indicated that miR-29a and miR-34a may be important regulators of COL1A1 gene expression in CLs, possibly associated with an ultrastructural deterioration of these tissues during ageing. These miRNA: gene target interactions may be then responsible for ageing related pathophysiological processes in CLs through regulation of inflammatory related genes, Notch signalling, MAPK and p53 signalling.

## Acknowledgment

We would like to thank Mr. John Waters, Institute of Integrative Biology for his technical support with collection of C57BL/6 mice.

