## Supplementary Table 1 for "Age-related changes in microRNAs expression in cruciate ligaments of wild-stock house mice"

Supplementary Table 1: List of used microRNA primers as below

| **Product** | **Product number** | **Source** |
| --- | --- | --- |
| miRScript RT II | 218161 | Qiagen |
| miRScript SybrGreen | 218073 | Qiagen |
| RNU-6 qPCR primer | MS00033740 | Qiagen |
| miR-128 primer | MS00008582 | Qiagen |
| miR-455 primer | MS00009744 | Qiagen |
| miR-29a primer | MS00003262 | Qiagen |
| miR-143 primer | MI0000459 | Qiagen |
| miR-21 primer | MI0000077 | Qiagen |
| miR-34a primer | MI0000268 | Qiagen |
| miR-181a primer | MS00011263 | Qiagen |
| miR-181b primer | MS00006699 | Qiagen |
| miR-181c primer | MS00008841 | Qiagen |
| miR-181d primer | MS00031500 | Qiagen |
