## Supplementary Table 2 for "Age-related changes in microRNAs expression in cruciate ligaments of wild-stock house mice"

| **Gene** | **Forward primer** | **Reverse primer** |
| --- | --- | --- |
| GAPDH | GAGAGGCCCTATCCCAACTC | GTGGGTGCAGCGAACTTTAT |
| COL1A1 | TGACTGGAAGAGCGGAGAGT | CAGACGGCTGAGTAGGGAAC |
| COL3A1 | CTGTAACATGGAAACTGGGGAAA | CCATAGCTGAACTGAAAACCACC |
| COL5A1 | CCTGGCATCAACTTGTCCGATGG | GTGGTCACTGCGGCTGAGGAACTTC |
| COL5A2 | TGGGGACTGATGGTACACCT | GGATCACCCGATTGTCCTCG |
| COL12A1 | CCAGACGACCACGCTCAAT | TCTTCTCCATGACCGAAGTGG |
| PRELP | CCCACACCCAGATTTCCTCAG | TGGACAGTCAGGGAAGACAGA |

Supplementary Table 2: List of used mRNA primers sequences
